# Robustness testing and scalability of phosphate regulated promoters useful for two-stage autoinduction in *E. coli*

**DOI:** 10.1101/2020.01.26.920280

**Authors:** Eirik A. Moreb, Zhixia Ye, John P. Efromson, Jennifer N. Hennigan, Romel Menacho-Melgar, Michael D. Lynch

## Abstract

A key challenge in synthetic biology is the successful utilization of characterized parts, such as promoters, in different biological contexts. We report the robustness testing of a small library of *E. coli* PhoB regulated promoters that enable heterologous protein production in two-stage cultures. Expression levels were measured both in a rich Autoinduction Broth as well as a minimal mineral salts media. Media dependent differences were promoter dependent. 4 out of 16 promoters tested were identified to have tightly controlled expression which was also robust to media formulation. Improved promoter robustness led to more predictable scale up and consistent expression in instrumented bioreactors. This subset of PhoB activated promoters, useful for two-stage autoinduction, highlight the impact of the environment on the performance of biological parts, and the importance of robustness testing in synthetic biology.

**Highlights:**

- Characterization of the impact of media on promoter activity
- Identification of promoters robust to environmental variables
- Identification of promoters whose expression scale from microtiter plates to bioreactors

Graphical Abstract

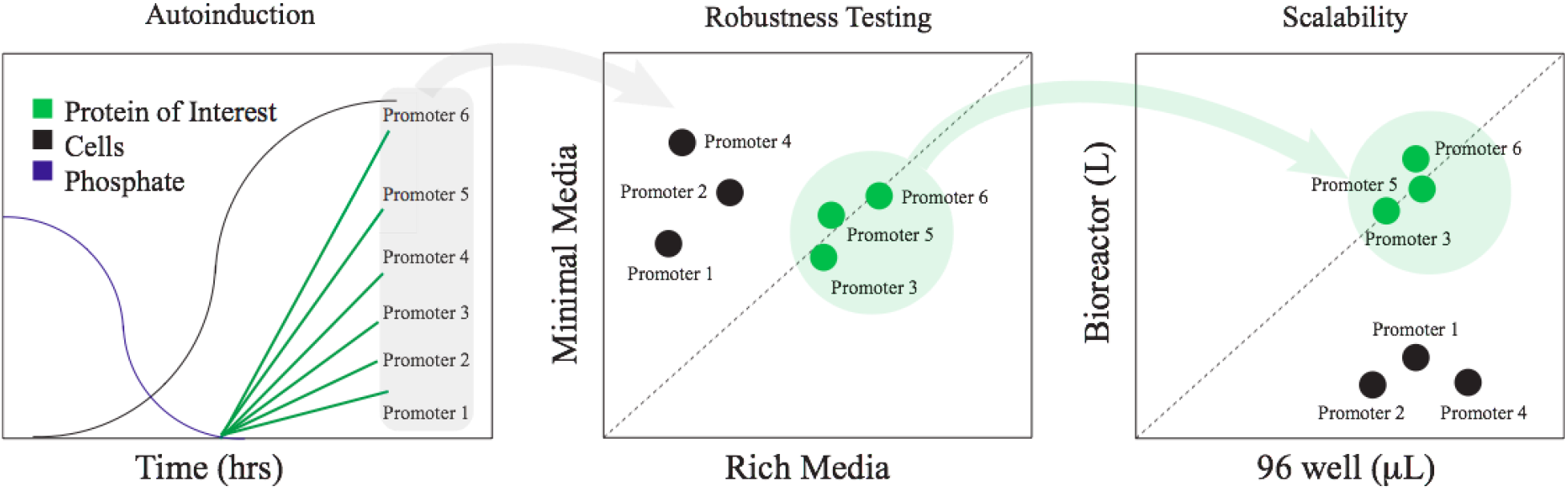

## Introduction

At the heart of synthetic biology, lies the characterization of standardized parts enabling the reuse of these components in novel engineered systems. ^1–3^ However, a critical challenge to synthetic biologists is that “part” performance is often dependent on the specific biological context, which for *E. coli,* can include the host strain chosen, additional genetic modifications and of course media as well as experimental protocols chosen for part characterization, as well as numerous additional factors.^4–7^ Even outputs as simple as protein expression can vary greatly as a function of context. However, despite being one of the simplest outputs, recombinant protein production is a mainstay in industrial biotechnology, with an estimated >$190 billion global annual market ^8,9^.

We have reported two-stage autoinduction of recombinant proteins in *E. coli* utilizing phosphate depletion, which can simplify expression protocols and lead to elevated protein titers. ^10,11^ This methodology utilizes engineered *E. coli* host strains as well as optimized Autoinduction Broth (AB) and a low phosphate inducible promoter derived from the *E. coli yibD* (*waaH*) gene promoter.^12,13^ This promoter led to similar expression levels in both rich AB as well as minimal mineral salts media in instrumented bioreactors. ^10,12–18^ Minimal media is often preferred for higher cell density commercial production, but richer media, specifically media enabling batch autoinduction, is easier to use in the laboratory and in higher throughput studies. We chose to investigate whether the robustness to media observed in the case of the modified *yibD* promoter is promoter specific or a general function of PhoB mediated activation and phosphate depletion. Additionally, we sought to characterize the environmental boundary conditions for the successful use of PhoB activated promoters for two-stage expression.

The Pho regulon is a well characterized response to phosphate limitation, utilizing the PhoRB two-component signal transduction system ^18–20^ PhoR responds to extracellular inorganic phosphate levels and activates PhoB, the transcriptional activator, responsible for gene regulation. According to the annotated EcoCyc database^3^ (https://ecocyc.org/), there are a total of 26 PhoB activated promoters (not including PhoB repressed promoters), as described in Supplemental Materials, Table S1. Many of these promoters have complex regulation with activators and/or repressors beyond PhoB including the use of several sigma factors. A set of 16 of these promoters (including the *yibD* (*waaH*) promoter) have minimal additional regulation (or predicted regulation) and utilize either sigma70 or sigmaS to initiate transcription. We further investigated the robustness of these 16 promoters and their potential use in applications, including two-stage protein expression.

## Results and Discussion

We first constructed a series of reporter vectors wherein each of the chosen 16 promoters was used to drive expression of GFPuv. Refer to Supplemental Tables S2 and S3 for a list of plasmids used in this study, and promoter sequences utilized. These reporter vectors, illustrated in Figure 1a, were initially evaluated for autoinduction in AB using host strain DLF_R002 (F-, λ-, Δ*(araD-araB)567, lacZ4787(del)(::rrnB-3), rph-1, Δ(rhaD-rhaB)568, hsdR51, ΔackA-pta, ΔpoxB, ΔpflB, ΔldhA, ΔiclR, ΔarcA)* ^10^ in the BioLector™ microreactor, enabling real time measurements of biomass and fluorescence (Figure 1b, 1c). A group of five promoters had high levels of GFPuv expression leading to protein titers > 1.75g/L. These included the *phoA, phoB, ugpB, ytfK* and *yibD* gene promoters, with the *phoA* and *ugpB* promoters having the highest expression levels. No significant differences in strain growth behavior was observed (Figure 1b).

**Figure 1:**
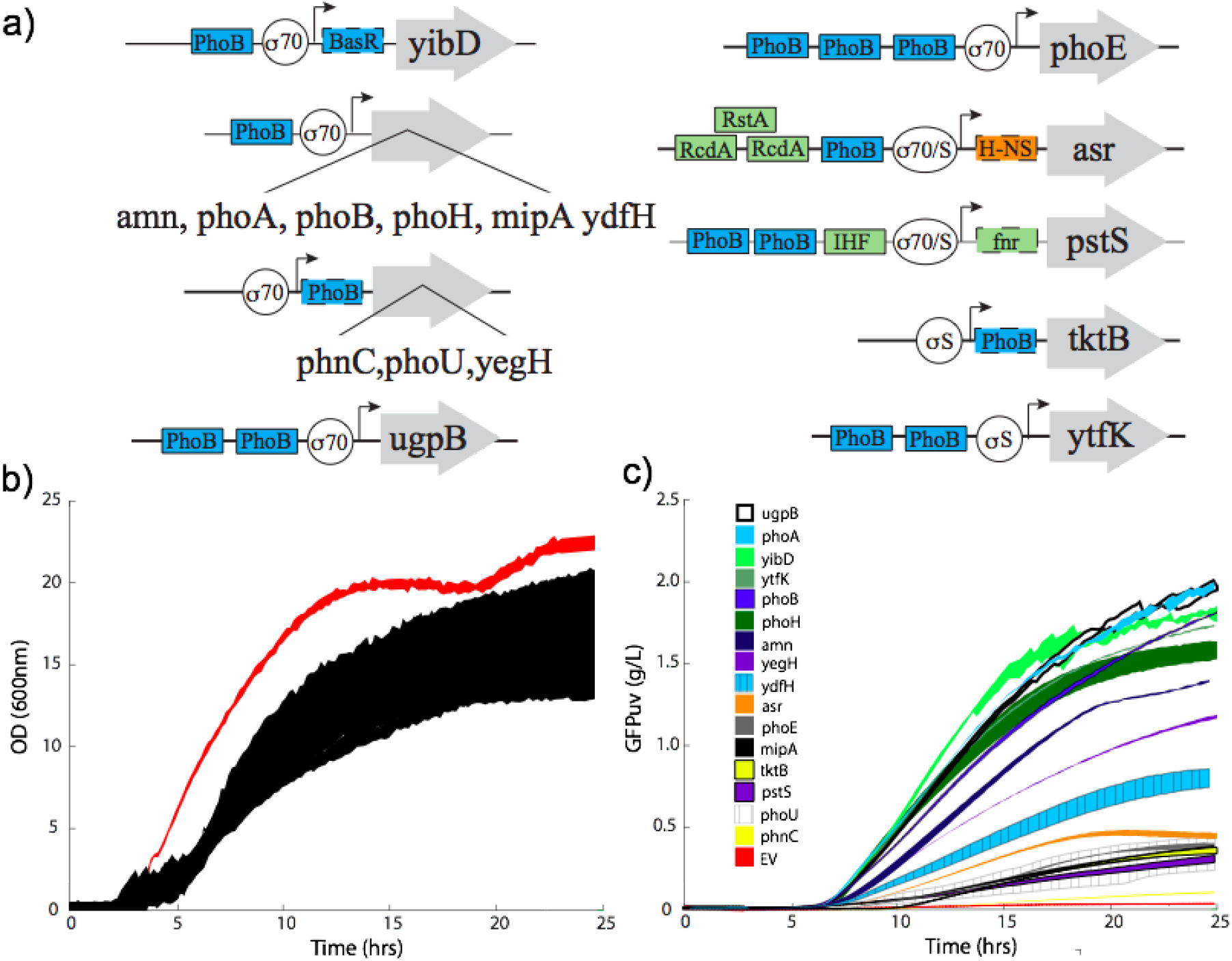
Characterization of PhoB activated promoters in autoinduction broth. a) The promoter architecture of a set of 16 previously characterized or predicted PhoB regulated promoters, that utilize sigma 70, sigma S or sigma 70 and sigma S for transcription. Known PhoB binding are shown as blue boxes. Additional activator (green) and repressor (orange) binding sites are shown. Boxes outlined with dashes have predicted but uncharacterized binding sites (Refer to Table 1 for References). b) Growth and c) autoinduction of GFPuv (c) by reporter vectors utilizing the PhoB regulated promoters. (b) red is empty vector (EV). Growth curves for all 16 reporter vectors, which overlap are shown in black. All curves are mean +/− one standard deviation from at least triplicate evaluations. All curves (except for the EV) were adjusted to have the same time of mid exponential point, which required a 3 hour adjustment at maximum. All vectors were evaluated in strain DLF_R002.

We next compared the activity of these promoters in AB to that in minimal mineral salts media. As minimal media fermentations in instrumented bioreactors are time consuming and not high throughput, we first adapted the microtiter plate based micro-fermentation protocol, as reported by Menacho-Melgar et al, to minimal media.^10^ This method relies on a seed (starter) culture in microtiter plates wherein minimal media is supplemented with small amounts of yeast extract and casamino acids. Cells are then harvested in mid-exponential phase after 16 hours of growth (5 < OD_600nm_ < 10), washed, and resuspended in minimal mineral salts media to deplete phosphate, while simultaneously enabling individual wells to be normalized to a similar optical density, in this case OD_600nm_=1. Results of microfermentations in both AB as well as minimal media are given in Figure 2. While the modified *yibD* promoter had similar expression levels in AB (~275mg/gCDW) and minimal mineral salts media (~225mg/gCDW), this was not true for most of the promoters characterized. For example, the *ytfK* promoter had a 60% reduction in expression in minimal media as compared to AB, while the *tktB* promoter had an almost 400% improvement in expression in minimal media. We next compared the behavior of the modified (truncated) *yibD* promoter ^10,11^ with the full length native promoter (Figure 2(e-j)). While both promoters had similar behavior in AB media, the full length promoter had a significant decrease (over 50% reduction) in expression in minimal media, presumably due to the presence of the extra NsrR repressor binding site or additional uncharacterized regulation.

**Figure 2:**
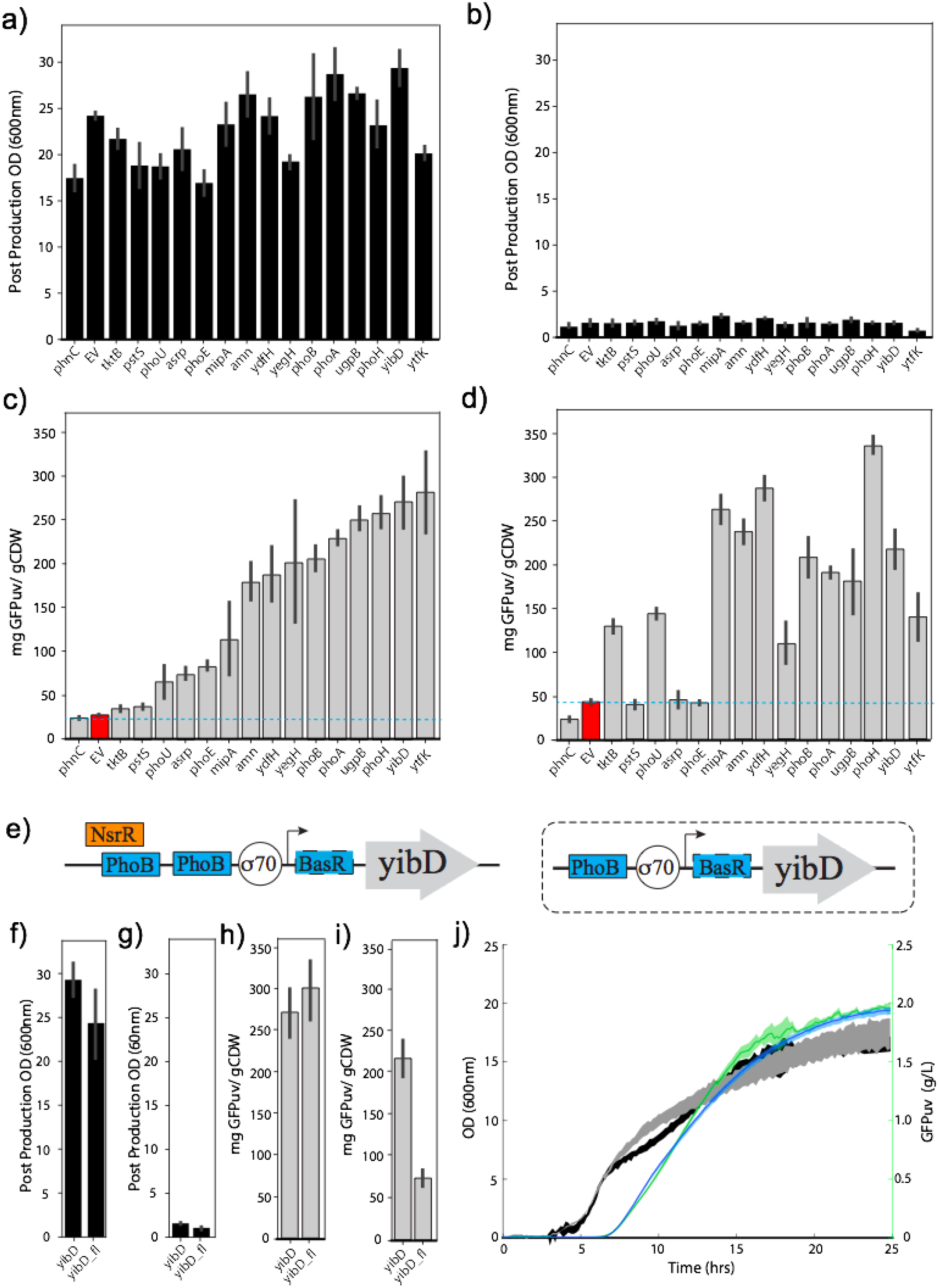
Comparison of PhoB regulated expression in AB with minimal media. All studies were carried out in 96 well microtiter plates. Final post production optical densities (OD_600nm_) for all strains either in a) Autoinduction broth or b) minimal media where cells were washed to deplete phosphate and normalized to OD_600nm_ =1. Specific GFP production (mgGFPuv/gCDW) at cell harvest for c) AB or d) minimal media. Empty vector control is highlighted in red. The highest specific production is indicated by a blue dashed line. (e through j) Comparison of the modified (truncated) and full length *yibD* promoter in AB and minimal media. e) The promoter architecture of the native yibD gene promoter (LEFT) and modified promoter truncated to remove a NsrR repressor binding site. Known PhoB binding are shown as blue boxes. BasR has a an uncharacterized binding site. Post Production optical density (OD_600nm_) for (f) AB and (g) minimal media. Specific GFPuv expression in (h) AB and (i) minimal media. j) Kinetics of growth and expression in AB in the BioLector ™. All studies used strain DLF_R002.

We next turned to evaluate the “leakiness” or basal activity for this group of promoters, including the insulated versions, in both minimal media and AB. In minimal media, we measured GFPuv fluorescence in mid-exponential cells growing in the seed cultures just prior to start of the production phase which occurs after cells are washed. In AB, data were taken from BioLector™ studies. As autoinduction occurs at the onset of the last cell doubling at OD_600nm_ from 7 to 10 (Figure 1c) we took GFPuv measurements early in exponential phase at OD_600nm_ = 3. Results are given in Figure 3 (a and b). In general, basal expression was higher in minimal media than AB, with several promoters having significant GFPuv expression prior to phosphate depletion. Specifically, the *amn, ydfH, yegH, ytfK, phoH* and *mipA* promoters had statistically significant basal expression levels above 20 mg/gCDW. These promoters were also the leakiest in AB, although the maximal basal expression observed was less than in minimal media. The promoters showing the tightest regulation in both minimal media and AB include the modified *yibD* promoter as well as the *asr, phoA, phoE* and *ugpB* promoters. A comparison of post-induction expression in the two different media is given in Figure 3c. While most promoters had significantly different expression in the two media, with most having improved expression in minimal media, there are several promoters with robust behavior and similar expression levels in both media. This includes the modified *yibD* promoter as well as the *phoA*, *phoB, ugpB, amn* and *pstS* promoters. Taken together, the set of tightly controlled promoters with reasonable expression and robustness to media include the modified *yibD, ugpB* and *phoA* promoters, with *phoB* having similar attributes but higher basal expression levels.

**Figure 3:**
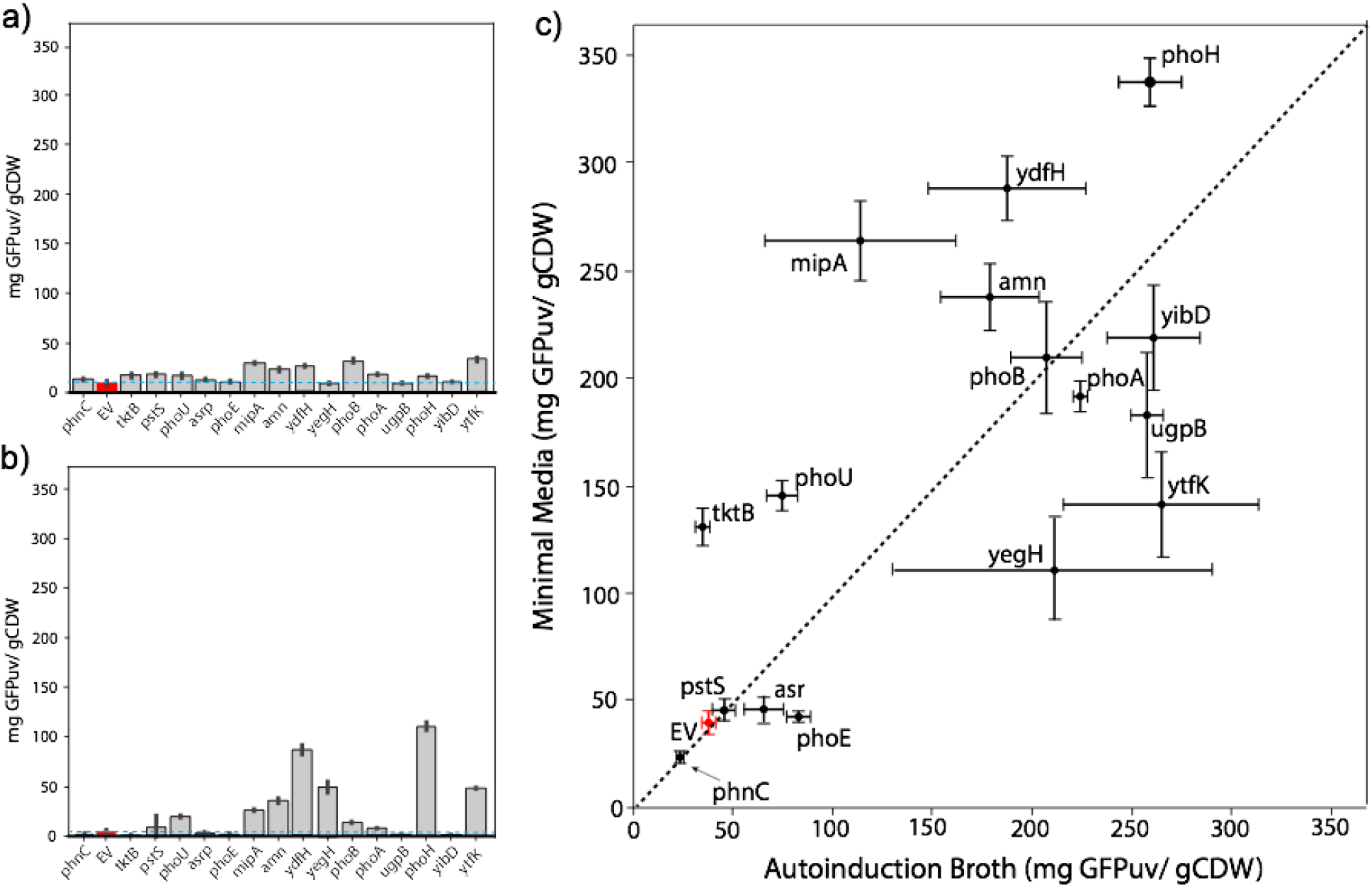
Environmental Dependence of PhoB regulated promoters. a) Leaky expression in AB. Specific GFPuv production (mg/gCDW) of promoters in AB in early exponential phase (OD_600nm_ = 3) b) Leaky expression in minimal media. Specific GFPuv production (mg/gCDW) of promoters in mid exponential phase in minimal media prior to wash and phosphate depletion. Empty vector is highlighted in red. The dashed blue line indicates the expression level of the empty vector control. c) Post induction specific expression in AB vs. Minimal Media. Empty vector is highlighted in red. Results are from 96 well microtiter fermentations, using DLF_R002 as a host strain.

Importantly, the *phoH* promoter with reasonable expression in AB, had a significant improvement in expression in minimal media, reaching the highest specific GFP levels (336 +/−22 mg/gCDW) on any promoter independent of media. This GFPuv level corresponds to ~ 66% of total cellular protein. *phoH* also had significant basal expression levels, which is consistent with a previous report that the *phoH* promoter region has at least 2 promoters, one inducible and one constitutive. ^21^ In fact, all of the promoters with higher specific expression levels in minimal media over AB (except for the *tktB* promoter), had leaky expression prior to phosphate depletion. These promoters, including the *phoH* promoter, may be of use where high expression levels are needed but tight control over expression is not a requirement.

The induction of these promoters was then characterized in response to oxygenation levels in cultures. We had previously reported that expression in AB using the modified yibD promoter was dependent on oxygenation.^10^ This necessitated optimizing culture volumes to enable adequate oxygenation in batch cultures. ^10^ We investigated the impact of oxygenation on promoter autoinduction in 96 well microtiter plates by changing culture volumes, with optimal aeration occurring at a fill volume of 100 μL and oxygen limiting conditions at fill volumes of 150 and 175 μL.^10^ Results are given in Figure 4. Unfortunately, most promoters had reduced expression under limited culture aeration as can be seen in Figure 4a. This impact is not solely a function of reduced growth rate leading to incomplete phosphate consumption in 24 hrs. Cultures extended to longer times, (48 hrs, Figure 5b) reach target optical densities without significant increases in GFPuv levels for the most tightly controlled promoters. Interestingly, a group of promoters with unwanted basal expression including the *mipA, phoU, and ydfH* promoter had robust behavior under limited aeration. The *phoE* and *tktB* promoters, which are tightly controlled yet with low levels of relative induction (Figure 2) also demonstrated robust behavior under limited aeration.

**Figure 4:**
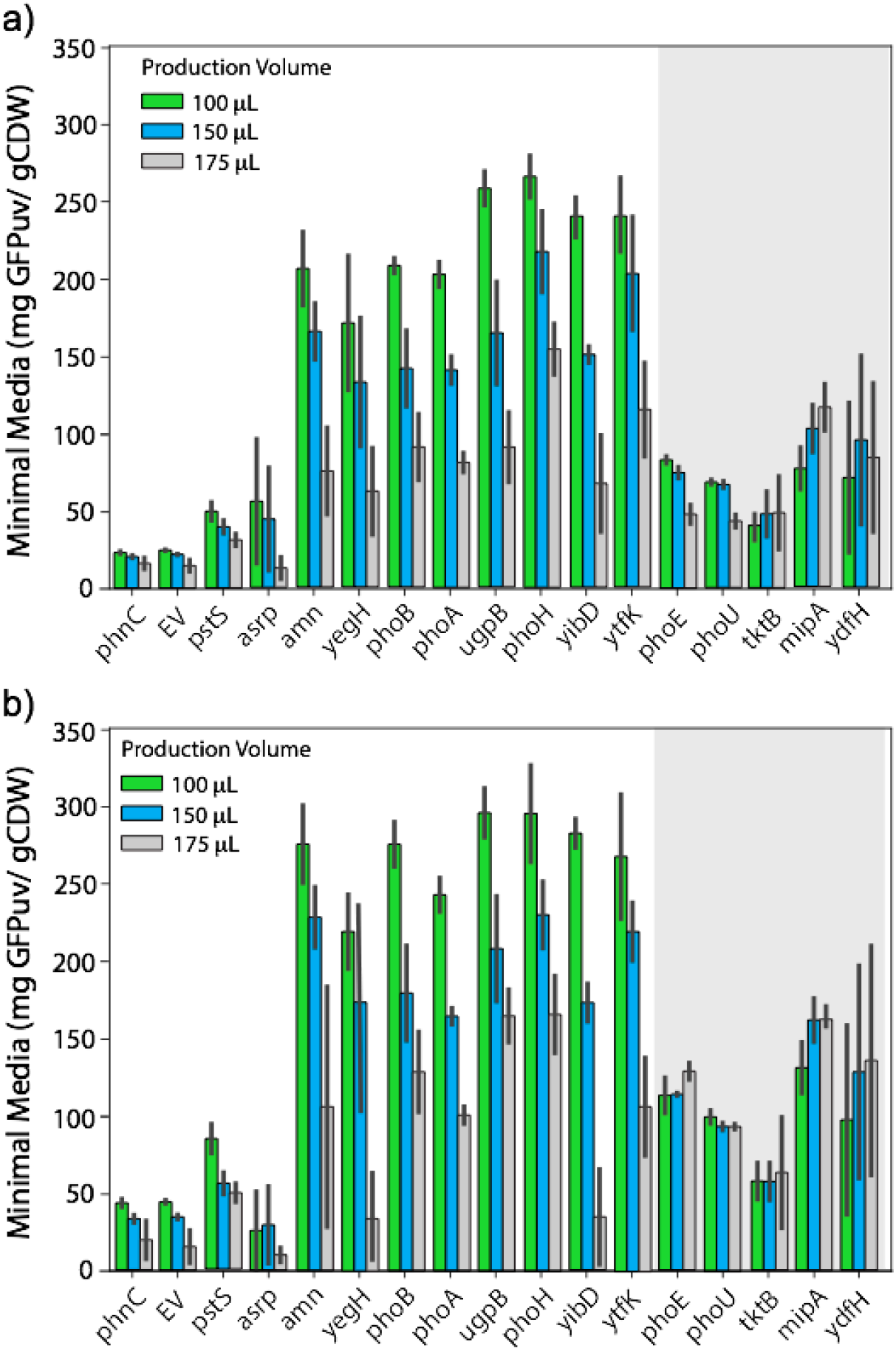
PhoB based autoinduction as a function of culture aeration (volume) in minimal media. a) Specific GFPuv expression 24 hours post wash as a function of 96 well microtiter plate volume. b) Specific GFPuv expression 48 hours post wash as a function of 96 well microtiter plate volume. Gray background indicates promoters with more robust behavior to oxygenation. Refer to Supplemental Figure S1 for biomass levels.

**Figure 5:**
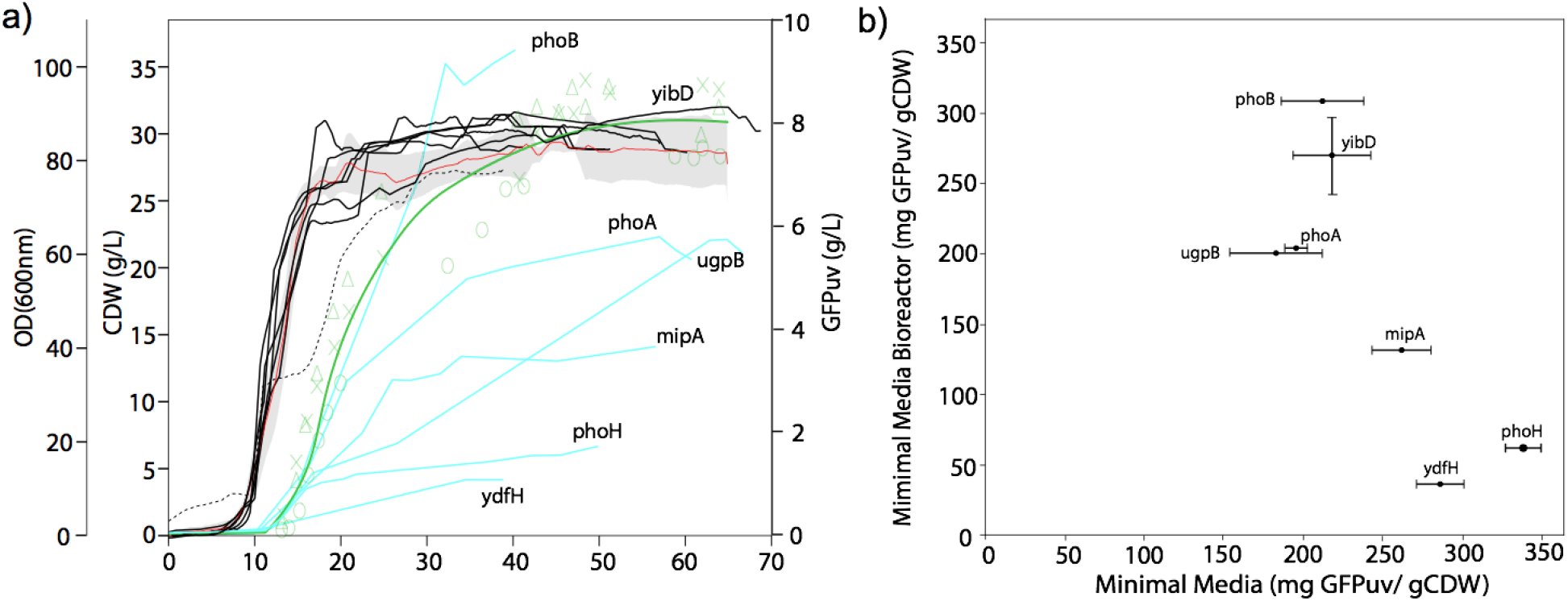
Scalability of a subset of PhoB activated promoters. a) Time course of growth and expression of GFPuv reporters in 30 gCDW/L fermentations in instrumented bioreactors. Control growth of strain DLF_R003 (DLF_R002, Δ*ompT*) with the *yibD* promoter is shown in red, with the error of three replicates indicated by the gray shaded region. GFP for this reporter is given by the green line which is the fit of three replicates (indicated by colored X’s circles and triangles).The *yibD* promoter data taken from Mencho Melgar et al. 10 Additional growth (black lines) and GFPuv expression (blue lines) profiles are plotted for strain DLF_R002 with the *mipA, phoA, phoB, phoH*, *ugpB,* and *ydfH* reporters. The *ydfH* reporter, which has a different growth profile is a dashed black line. b) Scalability for *mipA, phoA, phoB, phoH*, *ugpB, yibD* and *ydfH* reporters, plotting specific production in minimal media micro-fermentations (96 well plates) with specific production in instrumented bioreactors.

Lastly, we sought to evaluate the scalability of a subset of these PhoB activated promoters, by evaluation of autoinduction in minimal media in instrumented bioreactors. We had previously demonstrated the scalability of expression utilizing the modified *yibD* promoter, and compared these results to the bioreactor performance and expression levels using the group of tightly controlled and robust promoters (*phoA, ugpB,* and *phoB*) as well as a group of less robust promoters with increased basal expression, but elevated GFPuv titers in minimal media microfermentations including the *mipA, phoH* and *ydfH* promoters. Results are given in Figure 5. GFPuv expression levels scaled well when using the *yibD, phoA, phoB* and *ugpB* promoters, where specific expression levels were maintained from the small scale into bioreactors. In contrast, final specific GFPuv expression in bioreactors was greatly reduced in the case of the *mipA*, *phoH* and *ydfH* promoters, where significant basal expression was observed. These data support the following concept: Simply put, if a promoter or part is evaluated to have robustness to diverse environments at the small screening scale, the more likely this robustness will translate to the environmental variation that occurs during scale up, even if specific variation observed at larger scale cannot be testing precisely during screening.

Of 15 native PhoB promoters evaluated only 4 (*yibD*, *phoA*, *phoB* and *ugpB*) can be considered to meet the key specifications for extensible use: tight regulation, media robustness and scalability. These promoters (as well as insulated version) have utility in two-stage expression and production. ^22,23^ None of these 4 promoters were found to be robust to oxygen conditions, and optimal expression requires adequate culture aeration, limiting use in oxygen limited or anaerobic cultures.

More broadly, the characterization of a small library of PhoB activated promoters under diverse environmental conditions highlights the importance of environmental factors in synthetic biology. To be truly “plug and play”, biological parts need to be robust enough to environmental conditions that their performance is consistent in new target applications. Minimally, an understanding of the impact of key environmental variables on performance is needed to assess part suitability for a given application (ie, lab scale production or production in a bioreactor using minimal media).^24^ By analogy, in electrical engineering, electronic components (parts) always have design specifications which include boundary conditions for desired operation, whether these are key inputs (voltage, current) or environmental specifications including temperature, humidity, etc.^25,26^

Promoter, and part, robustness enables faster translation of performance from one context to another, such as from one media to another or across production scales. The lack of scalability of the *mipA*, *phoH* and *ydfH* promoters from minimal media in microfermentations to bioreactors, may indeed be due to protocol differences between the two experiments, including media differences. In the small scale, cells are grown with extra rich nutrients prior to wash and resuspension in minimal media. While significant effort could be made to optimize screening protocols that predict larger scale performance, these may likely be very different in the case of each promoter. Historically, the optimization on a case by case, or part by part, basis has lead to protocols and workflow that represent almost artisanal solutions which are not broadly applicable.^4–7^ Assessing, and potentially engineering, improved part robustness can lead to greater plug and play extensibility in synthetic biology. In many cases, this must also include a better understanding of context dependent activity. ^24,27,28^ However, experimental characterization of failure modes is often lacking but would enable evaluation of factors beyond just local part context (adjacent DNA).^29^ This is particularly important in biological systems where the unknown levels of regulation and control are almost universally encountered. Continued efforts to better characterize the impact of the context on part performance and the development of standards for part robustness will push synthetic biology forward as a true engineering discipline.

## Materials & Methods

### Reagents and Media

Unless otherwise stated, all materials and reagents were of the highest grade possible and purchased from Sigma (St. Louis, MO). Luria Broth, lennox formulation with lower salt was used for routine strain and plasmid propagation and construction and is referred to as LB below. AB, FGM10, FGM30, SM10 and SM10++ seed media were as described by Menacho-Melgar et al.^10^ Kanamycin was used at a final working concentration of 35 μg/mL. Yeast extract and MOPS (3-(N-morpholino)propanesulfonic acid) were obtained from Biobasic (Amherst, NY)., product numbers G0961 and MB0360, respectively. Enzyscreen 96 well microtiter sandwich plate covers (CR1596, Enzyscreen, The Netherlands) were used in micro-fermentations to reduce evaporative loss as high mixing speeds.

### Strains and Plasmids

*E. coli* strains DLF_R002 and DLF_R003 were constructed as previously described.^10^ Reporter plasmids were constructed in the pSMART-HC-Kan (Lucigen, WI) backbone. Promoter sequences are available in the Supplemental Materials. Phosphate promoters and pHCKan-yibDp-GFPuv backbone ^10^ were PCR amplified from *E. coli* genomic DNA using primers listed in Table S4. Reporter plasmids were assembled by Gibson Assembly using promoter PCR product and pHCKan-yibDp-GFPuv PCR product following manufacturer’s protocol (New England Biolabs, Ipswhich, MA). All plasmid sequences were confirmed by DNA sequencing (Genewiz, NC or Eurofins, KY). Sequences and maps are available with Addgene. Refer to Table 2 for Addgene numbers.

### Growth and Expression Analysis

Fluorescence measurements, GFPuv quantification, BioLector™, and microtiter plate expression using AB were all performed as previously reported. ^10^ Strain dry cell weight was correlated to OD600nm, using 1 OD600nm = 0.35 gCDW/L, according to Menacho Melgar et al. ^10^ Briefly, for microtiter plate expression using AB media ^10^ 96-well microtiter plates were filled with 100 μL of LB per well and then inoculated from frozen glycerol stocks. Plates were covered with sandwich covers (Model # CR1596, 96 well plates) and incubated at 37°C and 300 rpm for 16 hours. After 16 hours, 1 μL of the overnight culture was used to inoculate 100 μL of AB media. Plates were again covered and incubated under similar conditions for 24 hours. At 24 hours, GFPuv and OD were measured. For minimal media microfermentations, 5 μL of glycerol stocks were used to inoculate 96 well microtiter plates containing 150 μL per well of SM10++ (Supplemental Materials, Table S5). Plates were then incubated for 16 hours at 37°C and 300 rpm. After 16 hours, cells were washed to remove phosphate and inoculated into phosphate-free media, SM10 Production Media (Supplemental Materials, Table S6). The process for washing the cells was as follows: centrifuge the plates, discard the supernatant, fill with 150 μL of SM10 Production Media, centrifuge the plates again, discard the supernatant, and fill with 50 μL of SM10 Production Media. At this point, cells were resuspended and OD was measured. The median OD of the plate was then used to normalize the production plates to 1 OD in 150 μL volume. For example, if the median OD was 15, 10 μL of culture would be added to 140 μL of SM10 Production Media in new plates. The production plates were incubated at 37°C and 300 rpm for 24 hours. At 24 hours, GFPuv and OD were measured.

## Acknowledgements

We would like to acknowledge the following support: the NSF EAGER: #1445726 DARPA# HR0011-14-C-0075, ONR YIP #12043956, and DOE EERE grant #EE0007563, as well as the North Carolina Biotechnology Center 2018-BIG-6503. R. Menacho-Melgar and J.N. Hennigan were supported in part by the NIH Biotechnology Training Grant (T32GM008555).

## Author contributions

E.A. Moreb and Z. Ye contributed equally to this work. E. Moreb, J.N. Hennigan and R. Menacho-Melgar performed micro-fermentation and BioLector™ experiments. Z. Ye constructed plasmids and strains and performed biolector experiments. J.P. Efromson performed bioreactor studies. All authors wrote revised and edited the manuscript.

