## Supplementary Materials for "Robustness testing and scalability of phosphate regulated promoters useful for two-stage autoinduction in *E. coli*"

### Supplemental Materials

**Table S1:** A summary of known or predicted regulation of PhoB activated promoters

| | | | | |
|---|---|---|---|---|
| **Promoter** | **Sigma** | **Other Regulators** | **Regulators** | **Refs** |
| yibDp* | 70 | basR, nsrR | $PO_4^{-2}$ , NO | [1–3] |
| amnp1 | 70 | | $PO_4^{-2}$ | [1,4] |
| mipAp | 70 | | $PO_4^{-2}$ | [5,6] |
| phoAp | 70 | | $PO_4^{-2}$ | [2,7–9] |
| phoBp | 70 | | $PO_4^{-2}$ | [2,5,7,8,10] |
| phoEp | 70 | | $PO_4^{-2}$ | [2,11] |
| phoHp1 | 70 | | $PO_4^{-2}$ | [2,12] |
| phoUp | 70 | | $PO_4^{-2}$ | [2,13] |
| ugpBp1 | 70 | | $PO_4^{-2}$ | [14,15] |
| ydfHp | 70 | | $PO_4^{-2}$ | [5,6] |
| phnC | 70 | | $PO_4^{-2}$ | [16–18] |
| yegH | 70 | | $PO_4^{-2}$ | [5,6] |
| asrp | 70, S | rcdA, rstA, H-NS | $PO_4^{-2}$ , Stress , Stationary Phase | [19,20] |
| pstSp | 70, S | fnr | $PO_4^{-2}$, $O_2$ , Stationary Phase | [13,21–24] |
| tktB | S | | $PO_4^{-2}$ | [25] |
| ytfK | S | | $PO_4^{-2}$ | [1,2,26,27] |
| argP | 70 | argP | $PO_4^{-2}$ | [28,29] |
| cusR | 70 | cusR,hprR | $PO_4^{-2}$, $Cu^{+2}$, $H_2O_2$ | [2,5,30] |
| cusC | 70 | cusR,hprR | $PO_4^{-2}$, $Cu^{+2}$, $H_2O_2$ | [2,5,30] |
| hiuH | 70 | cusR,hprR | $PO_4^{-2}$, $Cu^{+2}$, $H_2O_2$ | [2,5,30,31] |
| psiE | 70 | CRP | $PO_4^{-2}$, cAMP | [2,32] |
| ompFp | 70, S | envY, fur, cpxR, IHF, crp, ompR, rstA, micF, rybB | $PO_4^{-2}$ , osmolarity, cAMP, pH , others | [33–41] |

| | | | | |
|---|---|---|---|---|
| gadX | S | gadE, H-NS, gadX, gadW, fnr, rutR | $PO_4^{-2}$, acid, $O_2$, Stationary Phase, pyrimidines | [42–46] |
| phoQp5 | 24 | | $PO_4^{-2}$, Heat, extracellular protein | [2,47] |
| adiC | 32 | gadRcs | $PO_4^{-2}$, Heat shock, acid | [2,47,48] |
| yhjCp | 54 | | $PO_4^{-2}$, Nitrogen Limitation | [2,5,49,50] |
| *the modified yibDp promoter is truncated to remove the NsrR repressor binding site. | | | | |

**Table S2:** Plasmids used in this study

| **Name** | **Promoter** | **Addgene #** | **Source** |
|---|---|---|---|
| pHCKan-yibDp-GFPuv | Modified yibDp | 127078 | [51] |
| pHCKan-yibDp_fl-GFPuv | Full length yibDp | 139083 | This study |
| pHCKan-amnp-GFPuv | amnp1 | 139070 | This study |
| pHCKan-mipAp-GFPuv | mipAp | 139071 | This study |
| pHCKan-phoAp-GFPuv | phoAp | 139072 | This study |
| pHCKan-phoBp-GFPuv | phoBp | 139073 | This study |
| pHCKan-phoEp-GFPuv | phoEp | 139074 | This study |
| pHCKan-phoHp-GFPuv | phoHp | 139075 | This study |
| pHCKan-phoUp-GFPuv | phoUp | 139076 | This study |
| pHCKan-pstSp-GFPuv | pstSp | 139077 | This study |
| pHCKan-ugpBp-GFPuv | ugpBp1 | 139078 | This study |
| pHCKan-ydfHp-GFPuv | ydfHp | 139079 | This study |
| pHCKan-phnCp-GFPuv | phnCp | 139080 | This study |
| pHCKan-tktBp-GFPuv | tktB | 139081 | This study |
| pHCKan-asrp-GFPuv | asrp | 139082 | This study |
| pHCKan-yegH-GFPuv | yegH | 139084 | This study |
| pHCKan-ytfK-GFPuv | ytfK | 139085 | This study |

**Table S3:** Phosphate inducible promoter sequences evaluated, the ribosomal binding site is underlined, and the start codon of the reporter gene (GFPuv) is shown in green.

| Promoter Name | Sequence |
|---|---|
| ugpBp | TCTTTCTGACACCTTACTATCTTACAAATGTAACAAAAAAGTTATTTTTCTGT<br>AATTCGAGCATGTCATGTTACCCCGCGAGCATAAAACGCGTGTGTAGGAGGA<br>TAATCTATG |
| modified yibDp | GTGCGTAATTGTGCTGATCTCTTATATAGCTGCTCTCATTATCTCTCTACCCTG<br>AAGTGACTCTCTCACCTGTAAAAATAATATCTCACAGGCTTAATAGTTTCTTA<br>ATACAAAGCCTGTAAAACGTCAGGATAACTTCTGTGTAGGAGGATAATCTAT<br>G |
| yibDp_fl | TTTTATCTGAGGATAATCTGTTAAATATGTAAAATCCTGTCAGTGTAATAAAG<br>AGTTCGTAATTGTGCTGATCTCTTATATAGCTGCTCTCATTATCTCTCTACCCT<br>GAAGTGACTCTCTCACCTGTAAAAATAATATCTCACAGGCTTAATAGTTTCTT<br>AATACAAAGCCTGTAAAACGTCAGGATAACTTCTTGTAGGAGGATAATCTAT<br>G |
| phoAp | CGATTACGTAAAGAAGTTATTGAAGCATCCTCGTCAGTAAAAAGTTAATCTTT<br>TCAACAGCTGTCATAAAGTTGTCACGGCCGAGACTTATAGTCGCTTTGTTTTT<br>ATTTTTTAATGTATTTGTATGTAGGAGGATAATCTATGGCTAGCAAAGGAGA<br>AGAACTTTTCACATG |
| phoBp | GCCACGGAAATCAATAACCTGAAGATATGTGCGACGAGCTTTTCATAAATCT<br>GTCATAAATCTGACGCATAATGACGTCGCATTAATGATCGCAACCTATTTATT<br>GTGTAGGAGGATAATCTATGGCTAGCAAAGGAGAAGAACTTTTCACATG |
| amnp | AGACAGTCAACGCGCTTGATAGCCTGGCGAAGATCATCCGATCTTCGCCTTA<br>CACTTTTGTTTCACATTTCTGTGACATACTATCGGATGTGCGGTAATTGTATA<br>GGAGGATAATCTATG |
| ydfHp | GCTATGCCGGACTGAATGTCCACCGTCAGTAATTTTTATACCCGGCGTAACTG<br>CCGGGTTATTGCTTGTCACAAAAAAGTGGTAGACTCATGCAGTTAACTCACTT<br>GTAGGAGGATAATCTATG |
| mipAp | CATCCATAAATTTTGCATAATTAATGTAAAGACCAGGCTCGCCAGTAACGCT<br>AAATTCATTTGGCTGTAAGCGCGGTGTCATCCGCGTCAGGAAAATTAAACAG<br>TTACTTTAAAAAATGAAAACGTAAAAAGGTTGGGTTTCGATGTATTGACGGG<br>TAAACTTTGTCGCCCGCTAAACATTTGTTTGTGTAGGAGGATAATCTATG |
| phoHp | AATCCTGCTGAAAGCACACAGCTTTTTTCATCACTGTCATCACTCTGTCATCT<br>TTCCAGTAGAAACTAATGTCACTGAAATGGTGTTTTATAGTTAAATATAAGTA<br>AATATATTGTTGCAATAAATGCGAGATCTGTTGTACTTATTAAGTAGCAGCGG<br>AAGTTCTGTAGGAGGATAATCTATG |
| phoUp | ACCGAACTGAAGCAGGATTACACCGTGGTGATCGTCACCCACAACATGCAGC<br>AGGCTGCGCGTTGTTCCGACCACACGGCGTTTATGTACCTGGGCGAATTGATT<br>GAGTTCAGCAACACGGACGATCTGTTCACCATGTAGGAGGATAATCTATG |
| pstSp | AAGACTTTATCTCTCTGTCATAAAACTGTCATATTCCTTACATATAACTGTCA<br>CCTGTTTGTCCTATTTTGCTTCTCGTAGCCAACAAACAATGCTTTATGATGTA<br>GGAGGATAATCTATGGCTAGCAAAGGAGAAGAACTTTTCACATG |
| phoEp | AGCATGGCGTTTTGTTGCGCGGGATCAGCAAGCCTAGCGGCAGTTGTTTACG<br>CTTTTATTACAGATTTAATAAATTACCACATTTTAAGAATATTATTAATCTGT |

| | |
|---|---|
| | AATATATCTTTAACAATCTCAGGTTAAAAACTTTCCTGTTTTCAACGGGACTC TCCCGCTGTGTAGGAGGATAATCTATG |
| yegH | GTTATTACCCTCAAATAATATGAGATAAATAGTGCTGCAACATTGCATTTTTG CCCCGATTTATCCATGATCGAATTGTGACATTTGTCATACAACGTGTAGGAGG ATAATCTATG |
| ytfK | CTTCGGTAAAAAGATAATTCTGAATAATTGTAACCTTTAGGTAAAAAAAGTT ATACGCGGTGGAAACATTGCCCGGATAGTCTATAGTCACTAAGCATTAAAAT TTGCGCCTCATAATATGTAGGAGGATAATCTATG |

**Table S4: Oligonucleotides used in this study**

| Primer Name | Sequence |
|---|---|
| ugpBP-FOR | GTCCTGAATGATATCAAGCTTGAATTCGTT TCTTTCTGACACCTTACTATCTTACAAATG |
| ugpBP-REV | CTCCTTTGCTAGCCATAGATTATCCTCCTACACACGCGTTTTATGCTCGC |
| YibDp-FOR | GTCCTGAATGATATCAAGCTTGAATTCGTT GTGCGTAATTGTGCTGATCTC |
| YibDp-REV | CTCCTTTGCTAGCCATAGATTATCCTCCTACAAGAAGTTATCCTGACGTT TTACAGG |
| phoAp-FOR | GTCCTGAATGATATCAAGCTTGAATTCGTT CGATTACGTAAAGAAGTTATTGAAGCATC |
| phoAp-REV | CTCCTTTGCTAGCCATAGATTATCCTCCTACATACAAATACATTAAAAAA TAAAAACAAAGCGACTATAAG |
| phoBp-FOR | GTCCTGAATGATATCAAGCTTGAATTCGTT GCCACGGAAATCAATAACCTG |
| phoBp-REV | CTCCTTTGCTAGCCATAGATTATCCTCCTACAAATAAATAGGTTGCGATC ATTAATGCG |
| Amnp-FOR | GTCCTGAATGATATCAAGCTTGAATTCGTT AGACAGTCAACGCGCTTG |
| Amnp-REV | CTCCTTTGCTAGCCATAGATTATCCTCCTACAATACAATTACCGCACATC CGATAG |
| ydfHp-FOR | GTCCTGAATGATATCAAGCTTGAATTCGTT GCTATGCCGGACTGAATGTC |
| ydfHp-REV | CTCCTTTGCTAGCCATAGATTATCCTCCTACAAGTGAGTTAACTGCATGA GTCTAC |

| | |
|---|---|
| mipAp-FOR | GTCCTGAATGATATCAAGCTTGAATTCGTT CATCCATAAATTTTGCATAATTAATGTAAAGACC |
| mipAp-REV | CTCCTTTGCTAGCCATAGATTATCCTCCTACAAAACAAATGTTTAGCGGG CG |
| phoHp-FOR | GTCCTGAATGATATCAAGCTTGAATTCGTT AATCCTGCTGAAAGCACACAG |
| phoHp-REV | CTCCTTTGCTAGCCATAGATTATCCTCCTACAGAACTTCCGCTGCTACTT AATAAGTAC |
| yhjCp-FOR | GTCCTGAATGATATCAAGCTTGAATTCGTT CTACAGAGATGACGTGTAGAAAATAGTTAC |
| yhjCp-REV | CTCCTTTGCTAGCCATAGATTATCCTCCTACATTTCACAATGTTGTCATGC CG |
| phoUp-FOR | GTCCTGAATGATATCAAGCTTGAATTCGTT ACCGAACTGAAGCAGGATTAC |
| phoUp-REV | CTCCTTTGCTAGCCATAGATTATCCTCCTACATGGTGAACAGATCGTCCG |
| pstSp-FOR | GTCCTGAATGATATCAAGCTTGAATTCGTT AAGACTTTATCTCTCTGTCATAAAACTGTC |
| pstSp-REV | CTCCTTTGCTAGCCATAGATTATCCTCCTACATCATAAAGCATTGTTTGTT GGCTAC |
| phoEp-FOR | GTCCTGAATGATATCAAGCTTGAATTCGTT AGCATGGCGTTTTGTTGC |
| phoEp-REV | CTCCTTTGCTAGCCATAGATTATCCTCCTACACAGCGGGAGAGTCCC |
| pho-GFP-FOR | TGTAGGAGGATAATCTATGGCTAGC |
| pho-GFP-REV | AACGAATTCAAGCTTGATATCATTCAGG |

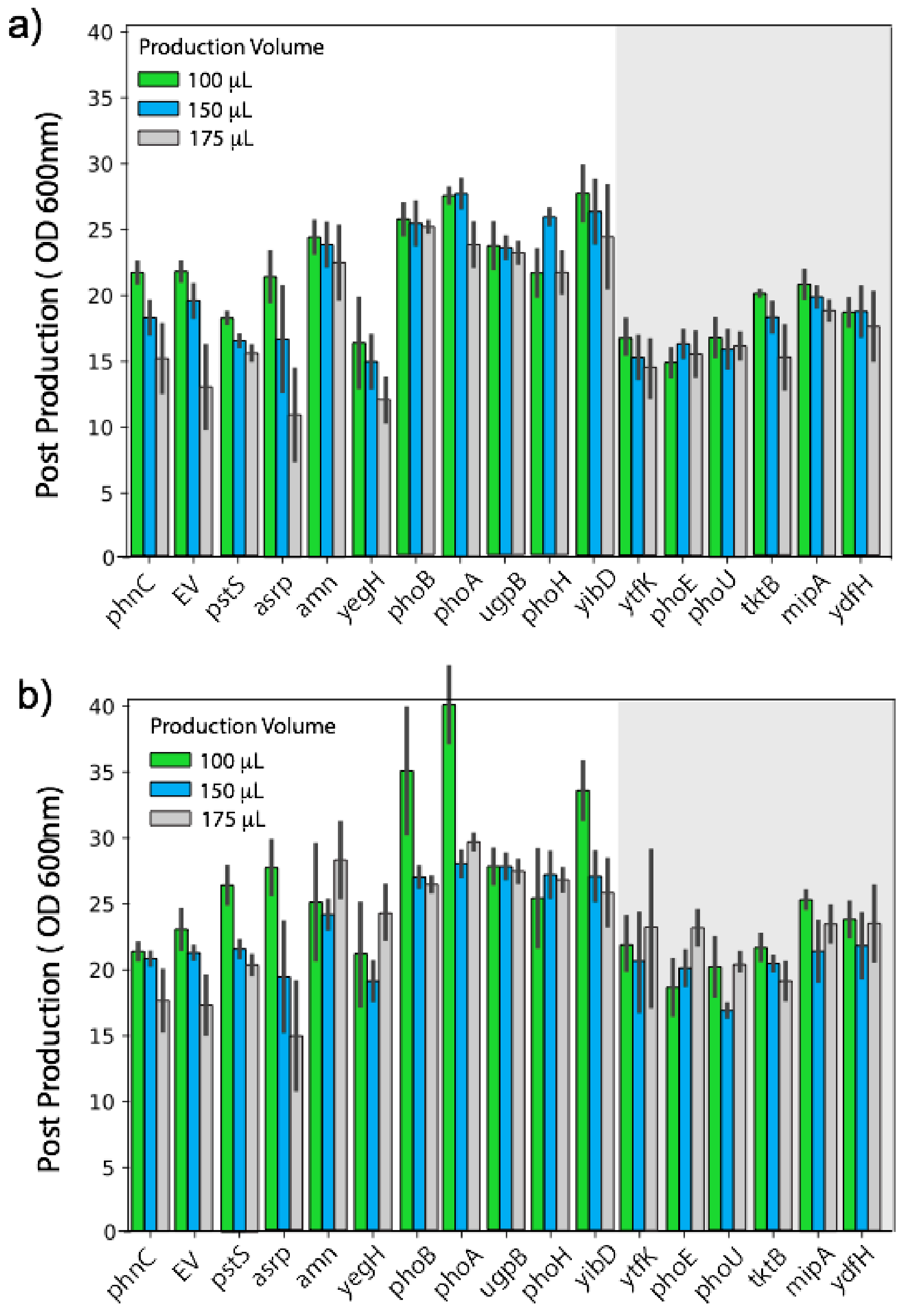


**Figure S1.** Strain growth as a function of culture aeration in minimal media. a) Post induction biomass levels ($OD_{600nm}$) as a function of 96 well microtiter plate volume, 24hours post wash. b) Post induction biomass levels ($OD_{600nm}$) as a function of 96 well microtiter plate volume, 48 hours post wash. Refer to Figure 5 in the main text for GFP expression.

**Media Stock Solutions**

- 10X concentrated Ammonium-Citrate 30 salts (1L) by mixing 30 g of $(NH_4)_2SO_4$ and 1.5 g Citric Acid in water with stirring, adjust pH to 7.5 with NaOH. Autoclave and store at room temperature (RT).
- 10X concentrated Ammonium-Citrate 90 salts (1L) by mixing 90 g of $(NH_4)_2SO_4$ and 2.5 g Citric Acid in water with stirring, adjust pH to 7.5 with NaOH. Autoclave and store at RT.
- 1 M Potassium 3-(N-morpholino) propanesulfonic Acid (MOPS), adjust to pH 7.4 with KOH. Filter sterilize (0.2 µm) and store at RT.
- 0.5 M potassium phosphate buffer, pH 6.8 by mixing 248.5 mL of 1.0 M $K_2HPO_4$ and 251.5 mL of 1.0 M $KH_2PO_4$ and adjust to a final volume of 1000 mL with ultrapure water. Filter sterilize (0.2 µm) and store at RT.
- 2 M $MgSO_4$ and 10 mM $CaSO_4$ solutions. Filter sterilize (0.2 µm) and store at RT.
- 50 g/L solution of thiamine-HCl. Filter sterilize (0.2 µm) and store at 4°C.
- 500 g/L solution of glucose, dissolving by stirring with heat. Cool, filter sterilize (0.2 µm), and store at RT.
- 100 g/L yeast extract, autoclave, and store at RT.
- 100 g/L casamino acid, autoclave, and store at RT.
- 500X Trace Metal Stock: Prepare a solution of micronutrients in 1000 mL of water containing 10 mL of concentrated $H_2SO_4$. 0.6 g $CoSO_4 \cdot 7H_20$, 5.0 g $CuSO_4 \cdot 5H_20$, 0.6 g $ZnSO_4 \cdot 7H_20$, 0.2 g $Na_2MoO_4 \cdot 2H_2O$, 0.1 g $H_3BO_3$, and 0.3 g $MnSO_4 \cdot H_2O$. Filter sterilize (0.2 µm) and store at RT in the dark.
- Prepare a fresh solution of 40 mM ferric sulfate heptahydrate in water, filter sterilize (0.2 µm) before preparing media each time.

Prepare the final working medium by aseptically mixing stock solutions based on the following tables in the order written to minimize precipitation, then filter sterilize (with a 0.2 µm filter).

**Table S6:** ***SM10++ Media, pH 6.8:***

| Ingredient | Concentration Stock | Volume in 1 L (mL) | Final Concentration |
|---|---|---|---|
| Ammonium-Citrate 90 Salts, pH 7.5 | 10 X | 100.0 | 1 X |
| Phosphate Buffer, pH 6.8 | 500 mM | 10.0 | 5.00 mM |

| | | | |
|---|---|---|---|
| Trace Metals | 500 X | 4.0 | 2 X |
| Fe (II) Sulfate | 40 mM | 4.0 | 0.16 mM |
| $MgSO_4$ | 2 M | 1.25 | 2.50 mM |
| $CaSO_4$ | 10 mM | 6.25 | 0.06 mM |
| Glucose | 500 g/L | 90.0 | 45 .0g/L |
| MOPS | 1 M | 200.0 | 200 mM |
| Thiamine-HCl | 50 g/L | 0.2 | 0.01 g/L |
| Yeast Extract | 100 g/L | 25.0 | 3.5 g/L |
| Casamino Acids | 100 g/L | 25.0 | 2.5 g/L |

**Table S7:** ***<u>SM10 Production Media (no phosphate), pH 6.8:</u>***

| Ingredient | Concentration Stock | Volume in 1 L (mL) | Final Concentration |
|---|---|---|---|
| Ammonium-Citrate 90 Salts, pH 7.5 | 10 X | 100.0 | 1 X |
| Phosphate Buffer, pH 6.8 | 500 mM | 0 | 0 |
| Trace Metals | 500 X | 4.0 | 2 X |
| Fe (II) Sulfate | 40 mM | 4.0 | 0.16 mM |
| $MgSO_4$ | 2 M | 1.25 | 2.50 mM |
| $CaSO_4$ | 10 mM | 6.25 | 0.06 mM |
| Glucose | 500 g/L | 90.0 | 45 .0g/L |
| MOPS | 1 M | 200.0 | 200 mM |
| Thiamine-HCl | 50 g/L | 0.2 | 0.01 g/L |
| Yeast Extract | 100 g/L | 0 | 0 |
| Casamino Acids | 100 g/L | 0 | 0 |

(5) Yang, C.; Huang, T.-W.; Wen, S.-Y.; Chang, C.-Y.; Tsai, S.-F.; Wu, W.-F.; Chang, C.-H. Genome-Wide PhoB Binding and Gene Expression Profiles Reveal the Hierarchical Gene

Regulatory Network of Phosphate Starvation in Escherichia Coli. *PLoS One* **2012**, *7* (10), e47314.
(6) Mendoza-Vargas, A.; Olvera, L.; Olvera, M.; Grande, R.; Vega-Alvarado, L.; Taboada, B.; Jimenez-Jacinto, V.; Salgado, H.; Juárez, K.; Contreras-Moreira, B.; et al. Genome-Wide Identification of Transcription Start Sites, Promoters and Transcription Factor Binding Sites in E. Coli. *PLoS One* **2009**, *4* (10), e7526.
(7) Makino, K.; Shinagawa, H.; Amemura, M.; Nakata, A. Nucleotide Sequence of the phoB Gene, the Positive Regulatory Gene for the Phosphate Regulon of Escherichia Coli K-12. *J. Mol. Biol.* **1986**, *190* (1), 37–44.
(8) Uluşeker, C.; Torres-Bacete, J.; García, J. L.; Hanczyc, M. M.; Nogales, J.; Kahramanoğulları, O. Quantifying Dynamic Mechanisms of Auto-Regulation in Escherichia Coli with Synthetic Promoter in Response to Varying External Phosphate Levels. *Sci. Rep.* **2019**, *9* (1), 2076.
(9) Yamada, M.; Makino, K.; Amemura, M.; Shinagawa, H.; Nakata, A. Regulation of the Phosphate Regulon of Escherichia Coli: Analysis of Mutant phoB and phoR Genes Causing Different Phenotypes. *J. Bacteriol.* **1989**, *171* (10), 5601–5606.
(10) Wanner, B. L.; Chang, B. D. The phoBR Operon in Escherichia Coli K-12. *J. Bacteriol.* **1987**, *169* (12), 5569–5574.
(11) Tommassen, J.; Koster, M.; Overduin, P. Molecular Analysis of the Promoter Region of the Escherichia Coli K-12 phoE Gene. Identification of an Element, Upstream from the Promoter, Required for Efficient Expression of phoE Protein. *J. Mol. Biol.* **1987**, *198* (4), 633–641.
(12) Kim, S. K.; Makino, K.; Amemura, M.; Shinagawa, H.; Nakata, A. Molecular Analysis of the phoH Gene, Belonging to the Phosphate Regulon in Escherichia Coli. *Journal of Bacteriology*. 1993, pp 1316–1324. https://doi.org/10.1128/jb.175.5.1316-1324.1993.
(13) Spira, B.; Aguena, M.; de Castro Oliveira, J. V.; Yagil, E. Alternative Promoters in the Pst Operon of Escherichia Coli. *Mol. Genet. Genomics* **2010**, *284* (6), 489–498.
(14) Kasahara, M.; Makino, K.; Amemura, M.; Nakata, A.; Shinagawa, H. Dual Regulation of the Ugp Operon by Phosphate and Carbon Starvation at Two Interspaced Promoters. *J. Bacteriol.* **1991**, *173* (2), 549–558.
(15) Overduin, P.; Boos, W.; Tommassen, J. Nucleotide Sequence of the Ugp Genes of Escherichia Coli K-12: Homology to the Maltose System. *Mol. Microbiol.* **1988**, *2* (6), 767–775.
(16) Jiang, W.; Metcalf, W. W.; Lee, K. S.; Wanner, B. L. Molecular Cloning, Mapping, and Regulation of Pho Regulon Genes for Phosphonate Breakdown by the Phosphonatase Pathway of Salmonella Typhimurium LT2. *J. Bacteriol.* **1995**, *177* (22), 6411–6421.
(17) Makino, K.; Kim, S. K.; Shinagawa, H.; Amemura, M.; Nakata, A. Molecular Analysis of the Cryptic and Functional Phn Operons for Phosphonate Use in Escherichia Coli K-12. *J. Bacteriol.* **1991**, *173* (8), 2665–2672.
(18) Wanner, B. L.; Metcalf, W. W. Molecular Genetic Studies of a 10.9-Kb Operon in Escherichia Coli for Phosphonate Uptake and Biodegradation. *FEMS Microbiol. Lett.* **1992**, *100* (1-3), 133–139.
(19) Suziedeliené, E.; Suziedélis, K.; Garbenciūté, V.; Normark, S. The Acid-Inducible Asr Gene in Escherichia Coli: Transcriptional Control by the phoBR Operon. *J. Bacteriol.* **1999**, *181* (7), 2084–2093.
(20) Seputiene, V.; Suziedelis, K.; Normark, S.; Melefors, O.; Suziedeliene, E. Transcriptional Analysis of the Acid-Inducible Asr Gene in Enterobacteria. *Res. Microbiol.* **2004**, *155* (7), 535–542.
(21) Taschner, N. P.; Yagil, E.; Spira, B. The Effect of IHF on sigmaS Selectivity of the phoA and Pst Promoters of Escherichia Coli. *Arch. Microbiol.* **2006**, *185* (3), 234–237.
(22) Taschner, N. P.; Yagil, E.; Spira, B. A Differential Effect of sigmaS on the Expression of the PHO Regulon Genes of Escherichia Coli. *Microbiology* **2004**, *150* (Pt 9), 2985–2992.
(23) Salmon, K.; Hung, S.-P.; Mekjian, K.; Baldi, P.; Hatfield, G. W.; Gunsalus, R. P. Global Gene Expression Profiling in Escherichia Coli K12. The Effects of Oxygen Availability and FNR. *J. Biol.*

*Chem.* **2003**, *278* (32), 29837–29855.
(24) Makino, K.; Shinagawa, H.; Amemura, M.; Kimura, S.; Nakata, A.; Ishihama, A. Regulation of the Phosphate Regulon of Escherichia Coli. Activation of pstS Transcription by PhoB Protein in Vitro. *J. Mol. Biol.* **1988**, *203* (1), 85–95.
(25) Jung, I. L.; Phyo, K. H.; Kim, I. G. RpoS-Mediated Growth-Dependent Expression of the Escherichia Coli Tkt Genes Encoding Transketolases Isoenzymes. *Curr. Microbiol.* **2005**, *50* (6), 314–318.
(26) Lacour, S.; Landini, P. SigmaS-Dependent Gene Expression at the Onset of Stationary Phase in Escherichia Coli: Function of sigmaS-Dependent Genes and Identification of Their Promoter Sequences. *J. Bacteriol.* **2004**, *186* (21), 7186–7195.
(27) Yoshida, Y.; Sugiyama, S.; Oyamada, T.; Yokoyama, K.; Kim, S.-K.; Makino, K. Identification of PhoB Binding Sites of the yibD and ytfK Promoter Regions in Escherichia Coli. *J. Microbiol.* **2011**, *49* (2), 285–289.
(28) Han, J. S.; Park, J. Y.; Lee, Y. S.; Thöny, B.; Hwang, D. S. PhoB-Dependent Transcriptional Activation of the iciA Gene during Starvation for Phosphate in Escherichia Coli. *Mol. Gen. Genet.* **1999**, *262* (3), 448–452.
(29) Celis, R. T. Repression and Activation of Arginine Transport Genes in Escherichia Coli K 12 by the ArgP Protein. *J. Mol. Biol.* **1999**, *294* (5), 1087–1095.
(30) Urano, H.; Umezawa, Y.; Yamamoto, K.; Ishihama, A.; Ogasawara, H. Cooperative Regulation of the Common Target Genes between $H_2O_2$-Sensing YedVW and $Cu^{2+}$-Sensing CusSR in Escherichia Coli. *Microbiology* **2015**, *161* (Pt 4), 729–738.
(31) Urano, H.; Yoshida, M.; Ogawa, A.; Yamamoto, K.; Ishihama, A.; Ogasawara, H. Cross-Regulation between Two Common Ancestral Response Regulators, HprR and CusR, in Escherichia Coli. *Microbiology* **2017**, *163* (2), 243–252.
(32) Kim, S. K.; Kimura, S.; Shinagawa, H.; Nakata, A.; Lee, K. S.; Wanner, B. L.; Makino, K. Dual Transcriptional Regulation of the Escherichia Coli Phosphate-Starvation-Inducible psiE Gene of the Phosphate Regulon by PhoB and the Cyclic AMP (cAMP)-cAMP Receptor Protein Complex. *J. Bacteriol.* **2000**, *182* (19), 5596–5599.
(33) Yoshida, T.; Qin, L.; Egger, L. A.; Inouye, M. Transcription Regulation of ompF and ompC by a Single Transcription Factor, OmpR. *J. Biol. Chem.* **2006**, *281* (25), 17114–17123.
(34) Tsung, K.; Brissette, R. E.; Inouye, M. Identification of the DNA-Binding Domain of the OmpR Protein Required for Transcriptional Activation of the ompF and ompC Genes of Escherichia Coli by in Vivo DNA Footprinting. *J. Biol. Chem.* **1989**, *264* (17), 10104–10109.
(35) Schmidt, M.; Zheng, P.; Delihas, N. Secondary Structures of Escherichia Coli Antisense micF RNA, the 5'-End of the Target ompF mRNA, and the RNA/RNA Duplex. *Biochemistry* **1995**, *34* (11), 3621–3631.
(36) Urban, J. H.; Vogel, J. Translational Control and Target Recognition by Escherichia Coli Small RNAs in Vivo. *Nucleic Acids Res.* **2007**, *35* (3), 1018–1037.
(37) Rampersaud, A.; Norioka, S.; Inouye, M. Characterization of OmpR Binding Sequences in the Upstream Region of the ompF Promoter Essential for Transcriptional Activation. *J. Biol. Chem.* **1989**, *264* (31), 18693–18700.
(38) Raivio, T. L.; Leblanc, S. K. D.; Price, N. L. The Escherichia Coli Cpx Envelope Stress Response Regulates Genes of Diverse Function That Impact Antibiotic Resistance and Membrane Integrity. *J. Bacteriol.* **2013**, *195* (12), 2755–2767.
(39) Batchelor, E.; Walthers, D.; Kenney, L. J.; Goulian, M. The Escherichia Coli CpxA-CpxR Envelope Stress Response System Regulates Expression of the Porins ompF and ompC. *J. Bacteriol.* **2005**, *187* (16), 5723–5731.
(40) Chubiz, L. M.; Rao, C. V. Role of the Mar-Sox-Rob Regulon in Regulating Outer Membrane Porin

Expression. *J. Bacteriol.* **2011**, *193* (9), 2252–2260.
(41) Andersen, J.; Delihas, N. micF RNA Binds to the 5’ End of ompF mRNA and to a Protein from Escherichia Coli. *Biochemistry* **1990**, *29* (39), 9249–9256.
(42) Shimada, T.; Hirao, K.; Kori, A.; Yamamoto, K.; Ishihama, A. RutR Is the Uracil/thymine-Sensing Master Regulator of a Set of Genes for Synthesis and Degradation of Pyrimidines. *Molecular Microbiology*. 2007, pp 744–757. https://doi.org/10.1111/j.1365-2958.2007.05954.x.
(43) Tramonti, A.; De Canio, M.; De Biase, D. GadX/GadW-Dependent Regulation of the Escherichia Coli Acid Fitness Island: Transcriptional Control at the gadY-gadW Divergent Promoters and Identification of Four Novel 42 Bp GadX/GadW-Specific Binding Sites. *Mol. Microbiol.* **2008**, *70* (4), 965–982.
(44) Constantinidou, C.; Hobman, J. L.; Griffiths, L.; Patel, M. D.; Penn, C. W.; Cole, J. A.; Overton, T. W. A Reassessment of the FNR Regulon and Transcriptomic Analysis of the Effects of Nitrate, Nitrite, NarXL, and NarQP as Escherichia Coli K12 Adapts from Aerobic to Anaerobic Growth. *J. Biol. Chem.* **2006**, *281* (8), 4802–4815.
(45) Giangrossi, M.; Zattoni, S.; Tramonti, A.; De Biase, D.; Falconi, M. Antagonistic Role of H-NS and GadX in the Regulation of the Glutamate Decarboxylase-Dependent Acid Resistance System in Escherichia Coli. *J. Biol. Chem.* **2005**, *280* (22), 21498–21505.
(46) Hommais, F.; Krin, E.; Laurent-Winter, C.; Soutourina, O.; Malpertuy, A.; Le Caer, J. P.; Danchin, A.; Bertin, P. Large-Scale Monitoring of Pleiotropic Regulation of Gene Expression by the Prokaryotic Nucleoid-Associated Protein, H-NS. *Mol. Microbiol.* **2001**, *40* (1), 20–36.
(47) Marzan, L. W.; Hasan, C. M. M.; Shimizu, K. Effect of Acidic Condition on the Metabolic Regulation of Escherichia Coli and Its phoB Mutant. *Arch. Microbiol.* **2013**, *195* (3), 161–171.
(48) Krin, E.; Danchin, A.; Soutourina, O. RcsB Plays a Central Role in H-NS-Dependent Regulation of Motility and Acid Stress Resistance in Escherichia Coli. *Res. Microbiol.* **2010**, *161* (5), 363–371.
(49) Barrios, H.; Valderrama, B.; Morett, E. Compilation and Analysis of sigma(54)-Dependent Promoter Sequences. *Nucleic Acids Res.* **1999**, *27* (22), 4305–4313.
(50) Zhao, K.; Liu, M.; Burgess, R. R. Promoter and Regulon Analysis of Nitrogen Assimilation Factor, sigma54, Reveal Alternative Strategy for E. Coli MG1655 Flagellar Biosynthesis. *Nucleic Acids Res.* **2010**, *38* (4), 1273–1283.
(51) Menacho-Melgar, R.; Ye, Z.; Moreb, E. A.; Yang, T.; Efromson, J. P.; Decker, J. S.; Lynch, M. D. Improved, Scalable, Two-Stage, Autoinduction of Recombinant Protein Expression in E. Coli Utilizing Phosphate Depletion. https://doi.org/10.1101/820787.
